# Moraxellaceae and *Moraxella* interact with the altered airway mycobiome in asthma

**DOI:** 10.1101/525113

**Authors:** Hai-yue Liu, Chun-xi Li, Zhen-yu Liang, Shi-yu Zhang, Wan-ying Yang, Yan-mei Ye, Yan-xia Lin, Rong-chang Chen, Hong-wei Zhou, Jin Su

## Abstract

Airway microbial-microbial interaction plays an important role in chronic airway inflammatory diseases such as asthma. *Moraxella* is widely regarded as a human respiratory tract pathogen. We aimed to investigate the interactions among *Moraxella,* Moraxellaceae (to which *Moraxella* belongs) and the airway microbiome in asthma. Induced sputum samples were obtained from 116 asthma patients and 29 healthy individuals, and the bacterial/fungal communities were profiled using 16S rRNA and ITS1 DNA gene sequencing. We found that asthma patients harboured significantly greater airway bacterial and fungal α-diversity than that of healthy individuals. Moraxellaceae, *Moraxella* and *Moraxella* otu19 (bacteria), and Schizophyllaceae, Polyporaceae, *Aspergillus, Schizophyllum* and *Candida* (fungi) were increased in the airway of asthma. Moreover, Moraxellaceae, Schizophyllaceae, Polyporaceae and *Candida* were positively associated with airway fungal α-diversity. Correlation networks revealed Moraxellaceae and *Moraxella* as microbial “hubs” in asthma that had significant negative connections with multiple bacterial communities, such as Leptotrichiaceae, Lachnospiraceae, Peptostreptococcaceae, Porphyromonadaceae, *Prevotella, Veillonella, Rothia* and *Leptotrichia*, but positive interactions with fungal communities such as Schizophyllaceae, Polyporaceae, *Candida* and *Meyerozyma*. Together, our finding revealed an altered microbiome and complex microbial-microbial interactions in the airway of asthma. Moraxellaceae and *Moraxella* showed significant interactions with the airway mycobiome, providing potential insights into the novel pathogenic mechanisms of asthma.

**IMPORTANCE:** With the advent of culture-independent techniques, growing evidence suggests that the airway microbiome is closely correlated with chronic respiratory diseases such as asthma. The complex microbial-microbial interaction exists in the airways of both healthy individuals and patients with respiratory diseases, which is of great significance for the pathogenesis and disease progression of asthma. In this study, we evaluated the airway dysbiosis in asthma patients, described the interaction between Moraxellaceae, *Moraxella* and airway bacterial/fungal communities, and it contributes to further understanding the pathogenic mechanisms of asthma.

## INTRODUCTION

Asthma is a common airway condition characterized by chronic airway inflammation that affects approximately 300 million people worldwide (1). Recently, new culture-independent techniques have revealed that human airways harbour unique microbial communities (including bacteria, fungi and viruses) that are closely correlated with chronic lung diseases such as asthma (2–4). The airway bacterial community in asthma patients differs significantly from that of healthy individuals, with increased bacterial diversity, and more Proteobacteria, especially *Moraxella*, and fewer Bacteroidetes (2).

Increasing evidence indicates that the airway bacterial microbiome plays an important role in asthma. In a study of more than 600 infants at 1 month of age, *Moraxella catarrhalis, Haemophilus influenzae* or *Streptococcus pneumoniae* cultured from the oropharynx was correlated with increased risks of later wheezing and asthma (5). Culture-independent techniques have suggested that airway bacteria may cause asthma and affect asthma activity (6). Several studies have indicated that airway bacteria are correlated with disease-related features and with the severity and therapeutic response of asthma (7–10). *Moraxella* is currently regarded as an important and common pathogen in the human respiratory tract and is associated with a range of diseases such as sinusitis, tracheitis, bronchitis and pneumonia in children and bronchitis, pneumonia and nosocomial infections in adults (11), as well as with exacerbation of asthma (12). In many asthma patients, *Moraxella catarrhalis* is a dominant pathogen in the airway bacterial community (10) and is one of the most frequently isolated bacteria from sputum cultures from patients with asthma exacerbations (13).

Recently, the presence of the fungal microbiome (mycobiome) in both healthy and abnormal lungs has been increasingly recognized, and this mycobiome may impact the clinical course of chronic pulmonary diseases such as asthma (14). One previous study identified differences in the airway mycobiome between asthma patients and healthy controls by using standard PCR techniques and pyrosequencing (15).

Airway bacterial-bacterial interactions are known to be associated with immune responses and airway inflammatory processes (16), thereby influencing the therapeutic response and clinical outcome of lung diseases (17). Recent findings have provided insights on the airway bacterial-fungal interaction, which may contribute to the decline in lung function and to disease progression (18) and is important in respiratory disease settings (19). However, fungal-bacterial interactions in asthma have not been studied extensively using culture-independent methods, and relatively little is known about the interaction among *Moraxella,* Moraxellaceae (to which *Moraxella* belongs) and the airway mycobiome in asthma.

Recognizing that complex airway microbial communities and microbial-microbial interactions play an increasingly important role in the development and progression of asthma (16, 19–21), we conducted a cross-sectional study to explore the alterations of Moraxellaceae and *Moraxella* in asthma airways as well as the interactions among Moraxellaceae, *Moraxella* and the airway mycobiome. We aimed to provide new insights on airway microbial characteristics, bacterial-fungal interactions, and the potential pathogenic mechanisms of asthma.

## RESULTS

### Clinical characteristics of the subjects

A total of 145 sputum samples were obtained from 116 asthma patients (55 stable asthma patients and 61 exacerbated asthma patients) and 29 healthy subjects. After the sequencing analysis, samples from 128 and 145 participants were analysed for bacterial and fungal community composition data, respectively. No significant differences were found in age, gender or smoking history between the healthy and asthmatic groups. Patients with stable asthma had more ICS usage than patients with exacerbated asthma (*P*=0.017) (Table 1).

**TABLE 1.** Clinical characteristics of the subjects.

| Parameters | Healthy<br>(n=29) | Asthma<br>(n=116) | *P* value | Stable asthma<br>(n=55) | Exacerbated asthma<br>(n=61) | *P* value |
| --- | --- | --- | --- | --- | --- | --- |
| Age (years) | 44.3±23.1 | 42.7±14.0 | .733 | 41.4±12.8 | 44.0±15.1 | .480 |
| Gender, n (male:female) | 21:8 | 62:54 | .065 | 28:27 | 34:27 | .603 |
| FEV1 (% pred) | NA | 87.9±18.8 | NA | 89.6±16.9 | 86.5±2.4 | .934 |
| FVC (% pred) | NA | 101.9±16.4 | NA | 102.3±15.6 | 101.5±17.2 | .506 |
| FEV1/FVC (%) | NA | 86.7±14.6 | NA | 87.9±12.6 | 85.7±16.2 | .771 |
| Smoking, n (yes:no) | 9:20 | 24:91 | .244 | 11:44 | 13:47 | .826 |
| ICS, n (yes:no) | NA | 78:38 | NA | 43:12 | 35:26 | .017* |

FEV1: forced expiratory volume in 1 second, FVC: forced vital capacity, ICS: inhaled corticosteroids.
The data are presented as n (chi-square test) or as the mean±standard deviation (independent t-test \* represents $P<.05$ ).

### Airway bacterial and fungal communities in asthma patients differed from those in healthy individuals

Compared with healthy individuals, asthma patients showed a significantly greater airway α-diversity (calculated using the Shannon index and PD_whole_tree index) (Fig. 1) and distinct differences in the community compositions (β-diversity) (Fig. S1A) of both bacterial and fungal communities.

**FIG 1.**
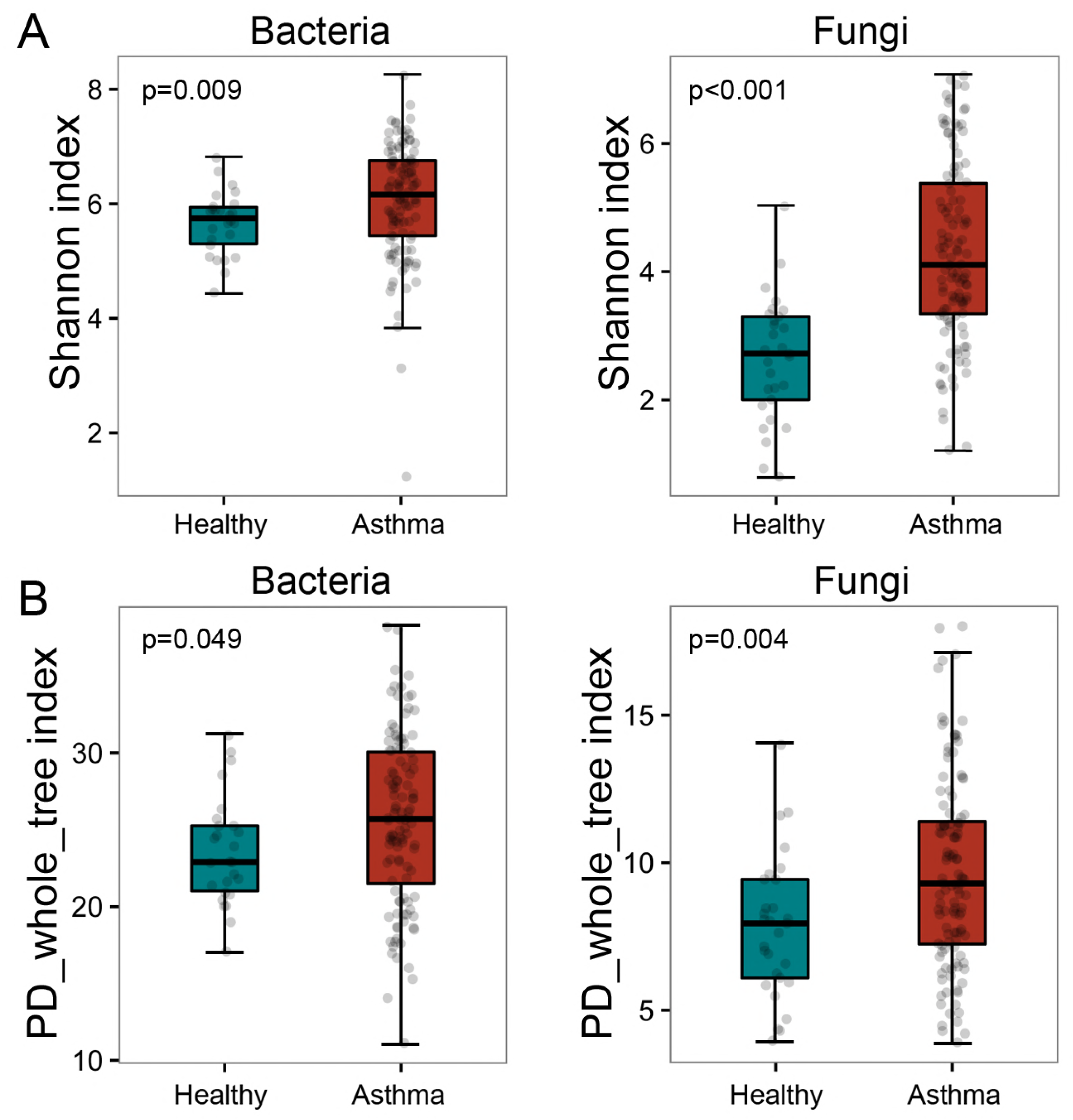
α-Diversity of the bacterial and fungal communities in healthy individuals and asthma patients. The Shannon index (A) indicates the evenness and richness diversity of the airway microbial community, and PD_whole_tree index (B) reflects the phylogenesis diversity of the airway microbial community. P values are presented using the Wilcoxon rank-sum test.

In addition, we compared the airway bacterial and fungal community compositions (relative abundance >1%) between asthma patients and healthy individuals (Table S1). Asthma patients harboured a slightly higher abundance of Proteobacteria (*P*=0.222) and an extremely increased abundance of Moraxellaceae, *Moraxella* and *Moraxella* otu19 in the airway than healthy individuals (Fig. 2). Among these selected bacterial and fungal communities, more than 70% of the fungal community differed significantly between asthma patients and healthy individuals, especially Ascomycota, Basidiomycota, Schizophyllaceae, Polyporaceae, *Aspergillus, Schizophyllum, Candida* and *Meyerozyma,* which were more prevalent in asthma patients, while only 20-45% of the bacterial community differed significantly between the two groups, suggesting that the airway fungal community had a more varied and complex dysbiosis than the bacterial community in asthma.

**FIG 2.**
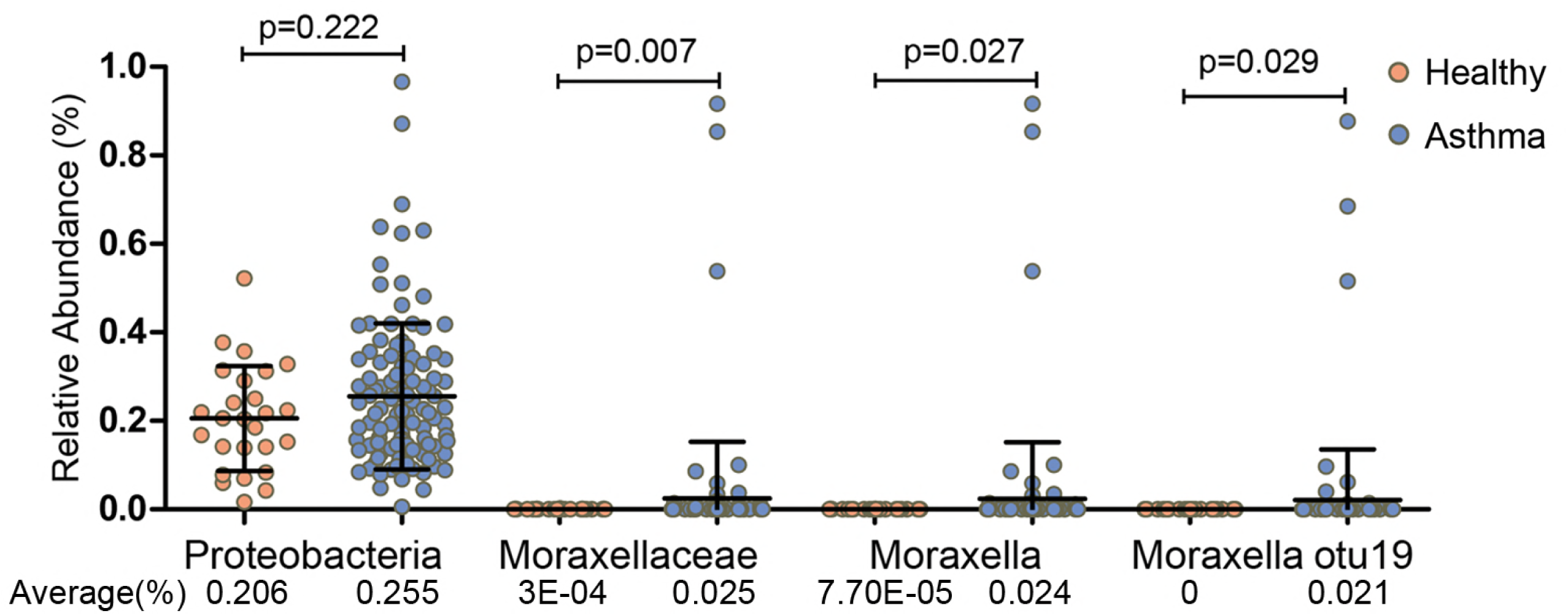
Comparison of the relative abundance of Proteobacteria, Moraxellaceae, *Moraxella* and *Moraxella* otu19 between healthy individuals and asthma patients. P values (Wilcoxon rank-sum test) and average relative abundances are presented.

Bronchial hyperresponsiveness in outpatients with asthma may affect the airway microbiome (22). Accordingly, we divided the asthmatic population into two subgroups—stable asthma and exacerbated asthma—according to disease activity. However, both the microbial α-diversity (Shannon index, bacteria: *P*=0.554, fungi: *P*=0.316, respectively) and β-diversity (bacteria: R^2^=0.009, *P*=0.269, fungi: R^2^=0.008, *P*=0.549, respectively) were similar, and no significant differences in either bacterial or fungal community composition were found between the two subgroups (Fig. S2).

Although not statistically significant, the phylum that exhibited the greatest increase in asthma was Proteobacteria. We therefore evaluated the overall families belonging to Proteobacteria. We found that the relative abundance of Moraxellaceae was significantly higher in asthma patients than in healthy individuals (80.5-fold, *P*=0.007) (Fig. 2). However, the abundances of other families (average abundance >1%), such as Neisseriaceae and Pasteurellaceae, were not significantly increased, and some were even decreased; for example, Burkholderiaceae was lower in asthma patients (*P*=0.041) (Fig. S1B). Focusing on the distinct difference in Moraxellaceae between asthma patients and healthy individuals, we further compared the different microbes at the genus level and OTU level. In agreement with the results for Moraxellaceae, *Moraxella* and *Moraxella* otu19 were significantly increased in asthma patients (314.7-fold, *P*=0.027; not detected in healthy individuals, *P*=0.029, respectively) (Fig. 2).

### Relationship among Moraxellaceae, *Moraxella* and airway bacterial/fungal diversity

As shown above, the airway microbial diversity in asthma patients differed from that in healthy individuals. Thus, we explored whether there was a relationship among the increased Moraxellaceae, *Moraxella* and airway bacterial/fungal α-diversity in asthma.

We first focused on the bacterial diversity and found no significant relationships among Proteobacteria, Moraxellaceae, *Moraxella, Moraxella* otu19 and bacterial diversity. However, a significantly positive correlation was found between Moraxellaceae and fungal diversity (Fig. 3). Next, we investigated whether the airway fungal community was related to fungal diversity and found that Basidiomycota, Schizophyllaceae, *Schizophyllum, Schizophyllum commune* otu6, *Aspergillus* otu5, Polyporaceae and *Candida* were positively correlated with fungal diversity (all *P* values<0.001) (Fig. S3). Our findings suggested that Moraxellaceae had a greater influence on the fungal community than the bacterial community in the asthma airway.

**FIG 3.**
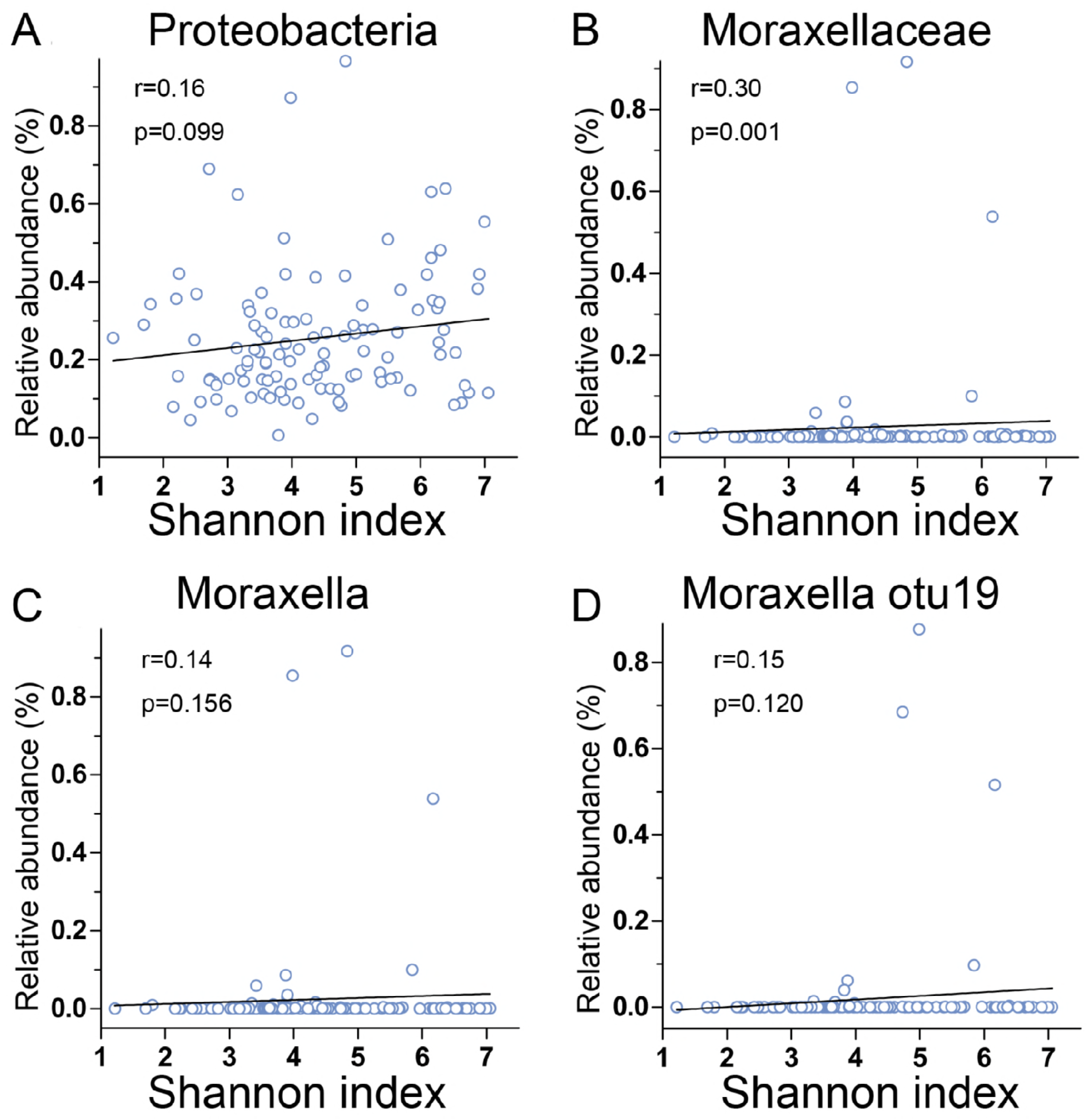
The relationships between Proteobacteria, Moraxellaceae, *Moraxella* and *Moraxella* otu19 and fungal α-diversity. R (cut-off of over 0.3) and P values are presented using Spearman rank correlation analysis.

### Interaction among Moraxellaceae, *Moraxella* and the airway mycobiome

To further explore the interaction among the increased Moraxellaceae, *Moraxella* and the airway microbiome in asthma, we used Spearman correlation analysis to build correlation networks (Fig. 4).

**FIG 4.**
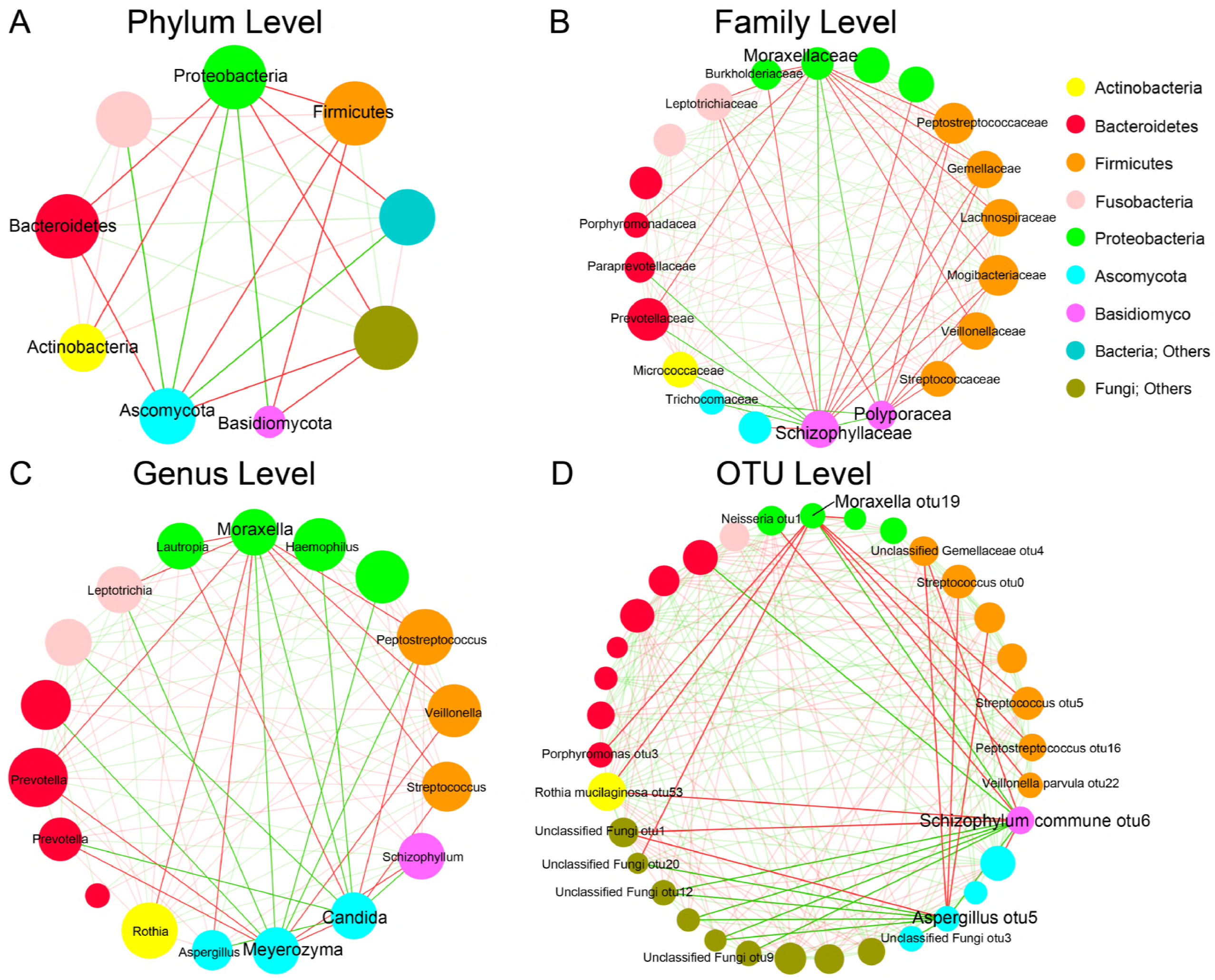
Correlation networks between airway microbial communities (microbial communities with a relative abundance of >1% are included) at the phylum (A), family (B), genus (C), and OTU (D) levels. Each node represents a microbial community, and its diameter is proportional to the degree of connectivity. The microbial communities are coloured by the phylum level. The red edges represent negative correlations, and the green edges represent positive correlations. The darker grey lines connect “hub” microbial communities that may play an important role in the asthmatic airway, and the lighter grey lines connect other airway microbial communities. Correlations were performed with Spearman rank correlation with a correlation cut-off P value of less than 0.05.

Examination of the microbial network revealed a few microbial “hubs” that were negatively connected with other portions of the airway microbiome. At the phylum level, Proteobacteria was significantly negatively associated with Firmicutes, Bacteroidetes and Actinobacteria but significantly positively correlated with Ascomycota and Basidiomycota. At the family level, Moraxellaceae showed 9 negative correlations with bacterial families, including with Veillonellaceae, Lachnospiraceae, Leptotrichiaceae, Prevotellaceae, Peptostreptococcaceae and Porphyromonadaceae, but 2 positive correlations with Schizophyllaceae and Polyporaceae. Moreover, Schizophyllaceae, Polyporaceae and Trichocomaceae exhibited strong positive correlations with each other and were negatively correlated with Leptotrichiaceae, Lachnospiraceae, Peptostreptococcaceae and Mogibacteriaceae. At the genus level, *Moraxella* was negatively associated with 7 bacterial genera (*Veillonella, Leptotrichia, Rothia, Streptococcus, Peptostreptococcus, Prevotella and Lautropia*) and positively associated with 2 fungal genera (*Candida* and *Meyerozyma). Candida* was positively related to *Aspergillus* and *Schizophyllum* but negatively linked to *Peptostreptococcus* and *Streptococcus. Meyerozyma* was negatively associated with *Prevotella* and *Veillonella.* At the OTU level, *Moraxella* otu19 was significantly negatively associated with *Veillonella parvula* otu22, *Streptococcus* otu5, *Peptostreptococcus* otu16 and Unclassified Fungi otu20 but significantly positively associated with *Schizophyllum commune* otu6 and Unclassified Fungi otu12. Additionally, *Schizophyllum commune* otu6 was negatively correlated with *Neisseria* otu1 and Unclassified Fungi otu1. Taken together, these results suggested complex and close interactions among Moraxellaceae, *Moraxella* and airway microbial communities (12 out of 21 communities and 10 out of 18 communities, respectively), especially the fungal community (Schizophyllaceae, Polyporaceae, *Candida* and *Meyerozyma*).

## DISCUSSION

Asthma is a chronic airway inflammatory disease associated with altered microbial communities in the airway, and these communities are closely related to airway inflammation. Recent studies have highlighted the importance of the complex microbial-microbial interactions in asthma (16, 18, 19). In addition to bacterial dysbiosis (23), we concurrently identified fungal dysbiosis characterized by alterations in diversity and composition in the asthma airway. We found that Moraxellaceae was positively associated with airway fungal diversity. Moraxellaceae and *Moraxella* showed positive interactions with Schizophyllaceae, Polyporaceae, *Candida* and *Meyerozyma* but negative interactions with Leptotrichiaceae, Lachnospiraceae, Burkholderiaceae, Mogibacteriaceae, Peptostreptococcaceae, Porphyromonadaceae, *Prevotella, Veillonella, Rothia, Leptotrichia* and *Streptococcus,* which were enriched in healthy individuals, indicating that Moraxellaceae and *Moraxella* interact with the altered airway mycobiome in asthma.

Our findings revealed significant differences in airway microbial diversity and community composition between asthma patients and healthy individuals. In agreement with previous studies, we found that asthma patients harboured a higher bacterial α-diversity (22, 23) and greatly increased proportions of Moraxellaceae and *Moraxella* in the airway compared with healthy individuals. Airway dysbiosis in asthma was evident not only in the bacterial community but also in the fungal community. Our study is the first to report that asthma patients have greater fungal α-diversity (1.6-fold greater Shannon index and 1.2-fold greater PD_whole_tree compared with healthy individuals) and more complicated fungal composition dysbiosis compared with bacterial dysbiosis (71.4-100% vs. 20-45%) in the airway.

*Moraxella* is widely recognized as a pathogen in the airway and is associated with several respiratory diseases such as asthma (11, 12). Multiple studies have reported altered airway microbial dysbiosis, with pathogenic Proteobacteria more frequently found in asthma (2, 22–25), and we found that this increase occurred mainly because of the increases in Moraxellaceae and *Moraxella,* which were 80.5-fold and 314.7-fold more abundant than in healthy individuals, respectively.

Previous studies demonstrated bacterial-bacterial interactions in chronic airway diseases such as asthma (16, 19). Our results showed unique microbial networks in asthma. Moraxellaceae and *Moraxella* were negatively correlated with 9 and 7 bacterial communities respectively, most of which included Leptotrichiaceae, Lachnospiraceae, Burkholderiaceae, Peptostreptococcaceae, Porphyromonadaceae, Mogibacteriaceae, *Prevotella, Veillonella, Rothia, Leptotrichia* and *Streptococcus.* These communities have been more prevalently found in healthy airways either in our findings or in previous studies (9, 23, 26, 27), suggesting that Moraxellaceae and *Moraxella* drive airway dysbiosis by inhibiting the commensal microbiome.

In the present study of airway microbial composition, Moraxellaceae had a greater influence on the α-diversity of fungi than bacteria. The dynamic interaction between bacteria and fungi may have dramatic effects on airway inflammatory processes (18, 28). The correlation network showed positive correlations connecting Moraxellaceae and *Moraxella* with Schizophyllaceae, Polyporaceae, *Candida* and *Meyerozyma.*

Additionally, *Schizophyllum,* which represents the vast majority of the family Schizophyllaceae, is reported as a respiratory pathogen. *Schizophyllum* appears to enhance both the severity and the exacerbation frequency of asthma (29); sensitization to *Schizophyllum* is an important risk factor for exacerbation frequency and a rapid decline in lung function in asthma (30). A previous study suggested that *Candida* may be pathogenically important in patients with asthma (31) and associated with severe exacerbations of asthma requiring hospital admission (32). We found that these pathogenic fungi were positively associated with Moraxellaceae and *Moraxella* but negatively associated with healthy enriched communities such as Leptotrichiaceae and *Leptotrichia*, which were more common in the asthma airway. These results suggest that Moraxellaceae and *Moraxella* have more significant effects on airway fungal communities than on bacterial communities. Moraxellaceae and *Moraxella* may influence the airway microbiome via interactions with the airway mycobiome, thus acting synergistically with pathogenic fungi to potentially drive the disease.

However, no significant differences were found between stable asthma and exacerbated asthma, suggesting that our outpatients with exacerbated asthma had disease that was extremely mild in severity without dyspnoea and easily controlled during exacerbation (33). Consequently, the microbial composition was similar between patients with exacerbated asthma and those with stable asthma.

Our study is strengthened by its relatively large sample size in China (9, 34) and by the combined exploration of airway bacterial and fungal communities. A limitation is the cross-sectional design, which cannot capture the dynamic changes in the airway microbiome during the progression of asthma.

In conclusion, we have revealed for the first time an increase in fungal α-diversity and more severe dysbiosis of fungi than of bacteria in the asthma airway. Importantly, our study explored the composition of Moraxellaceae and *Moraxella*, as well as the microbial-microbial interactions in the airway of asthma, and our results suggest that the interaction among Moraxellaceae, *Moraxella* and the airway mycobiome is important for further understanding airway dysbiosis in asthma. Our findings may provide a foundation for new insights on the pathogenic mechanisms of asthma. Further studies are needed to confirm the cause-effect relationship underlying the interaction of Moraxellaceae, *Moraxella* and the airway mycobiome in asthma.

## MATERIALS AND METHODS

### Study design and subjects

This study was approved by the ethics committee of Southern Medical University (Permit No. 2012-072). All subjects have provided written informed consent, in accordance with the Declaration of Helsinki. A total of 145 sputum samples were collected from 116 asthma patients and 29 healthy subjects enrolled at Nanfang Hospital, Southern Medical University (Guangzhou, China), between June 2015 and December 2016. Subjects provided clinical information including age, gender, forced expiratory volume in 1 second (FEV1), forced vital capacity (FVC), smoking history and use of inhaled corticosteroids (ICS).

Exacerbation of asthma is characterized by a progressive increase in symptoms of dyspnoea, cough, wheezing or chest tightness and a progressive decline of lung function, which represent changes in the patient’s usual state.

The inclusion criteria for asthma patients included age >15 years; initial diagnosis based on the Global Initiative for Asthma (GINA) guidelines (35); and a positive bronchodilator reversibility test result (FEV1 increased by >12% and 200 mL after inhaling 400 mg of salbutamol) or positive methacholine provocation test result. All subjects were free of clinical bacterial infection at the time of the study. Exclusion criteria included respiratory tract infection diagnosed by chest X-ray (every patient received a chest X-ray) within the past 4 weeks; the presence of any airway disease other than asthma; and peripheral white blood cell (WBC) counts outside the normal range.

### Sample collection, processing, DNA extraction and 16S rRNA/ITS1 gene amplification

For sputum induction and processing, the recommendations of the Task Force on Induced Sputum of the European Respiratory Society were used as the guidelines (36, 37). All samples were immediately stored at −80°C for subsequent DNA extraction after collection. The sputum samples were thawed under ventilation for 15 minutes, and genomic DNA extraction was performed using a total genomic DNA Nucleic Acid Extraction Kit (Bioeasy Technology, Inc., China) according to the manufacturer’s instructions.

The bacterial 16S rRNA genes and fungal ITS1 genes were amplified using the V4 region and ITS1 region of the genes, respectively. The amplicons were sequenced on an iTorrent sequencing platform. Detailed information on the DNA amplification and purification steps was described in our previous studies (38, 39).

### Statistical analysis

The raw sequence files for both the 16S rRNA V4 amplicons and fungal 18S–28S rRNA gene internally transcribed spacer regions ITS1 amplicons were processed by a barcoded Illumina paired-end sequencing (BIPES) pipeline (40). First, the barcode primers were trimmed and filtered if they contained ambiguous reads or mismatches in the primer regions according to the BIPES protocol. Then, we removed sequences that had more than one mismatch in the 40–70 bp regions. Next, we screened and removed chimaeras using UCHIME in de novo mode (41), and finally we generated the high-quality sequence reads of the 16S rRNA genes or ITS1 DNA genes.

We used the software pipeline “Quantitative Insights into Microbial Ecology” (QIIME) (1.9.1) to conduct subsequent analyses (42). Operational taxonomic units (OTUs) were clustered by using USEARCH with the default parameters and a threshold distance of .03. The Greengenes 13_8 database was used as a template file, and multiple alignments of QIIME-based sequences were performed using Python Nearest Alignment Space Termination (PyNAST) (43); representative sequences for the ITS1 gene were assigned with reference to the QIIME_ITS database (version information: sh_qiime_release_s_28.06.2017) (42); unclassified sequences were further defined by BLAST according to the closest shared taxonomic level. The representative 16S rRNA gene sequences were classified into specific taxa using the Ribosome Database Project (RDP) classifier (44). The sequences were deposited in the European Nucleotide Archive (ENA) under accession number PRJEB28853.

Alpha diversity metrics were analysed by the Shannon index, which indicates the evenness and richness of the airway microbial community, and PD_whole_tree, which reflects the phylogenesis of species. Beta diversity distances were calculated by principal coordinate analysis (PCoA) using unweighted_uniFrac distances, and statistical values were evaluated with the Adonis method. Differential features between groups were identified using a linear discriminant analysis (LDA) effect size (LEfSe) method with a threshold cut-off value for the logarithmic LDA score of 2.0. Because of the limitation of statistical power to detect uncommon OTUs, we selected the OTUs with an average relative abundance of >1% in any group. Statistical analyses of clinical parameters were performed using IBM SPSS version 20.0. The graphical representations were generated using GraphPad Prism 5, and the interaction network was constructed using Cytoscape (v3.6.0) (45).

## ACKNOWLEDGMENTS

This research was supported by the Guangzhou Healthcare Collaborative Innovation Major Project [201604020012], the National Key R&D Program of China [2017YFC1310601 and 2017YFC1310603], the National Natural Science Foundation of China [NSFC31570497] and the Open Project of the State Key Laboratory of Respiratory Disease [SKLRD2016OP014].

The authors’ responsibilities were as follows. H.L., C.L., H.Z., and J.S. designed the experiments. H.L., C.L., Z.L., S.Z., M.Y., M.Z., W.Y., and Y.L. collected samples and performed the experiments. H.L., C.L., and Z.L. analyzed the data. H.L., C.L., H.Z., and J.S. prepared the manuscript and had primary responsibility for final content. All authors read and approved the final manuscript. None of the authors reported a conflict of interest.

We thank the patients, clinical doctors and microbiology staff who supported this work. We thank American Journal Experts (AJE) for English language editing.

---

**FIG S1.**
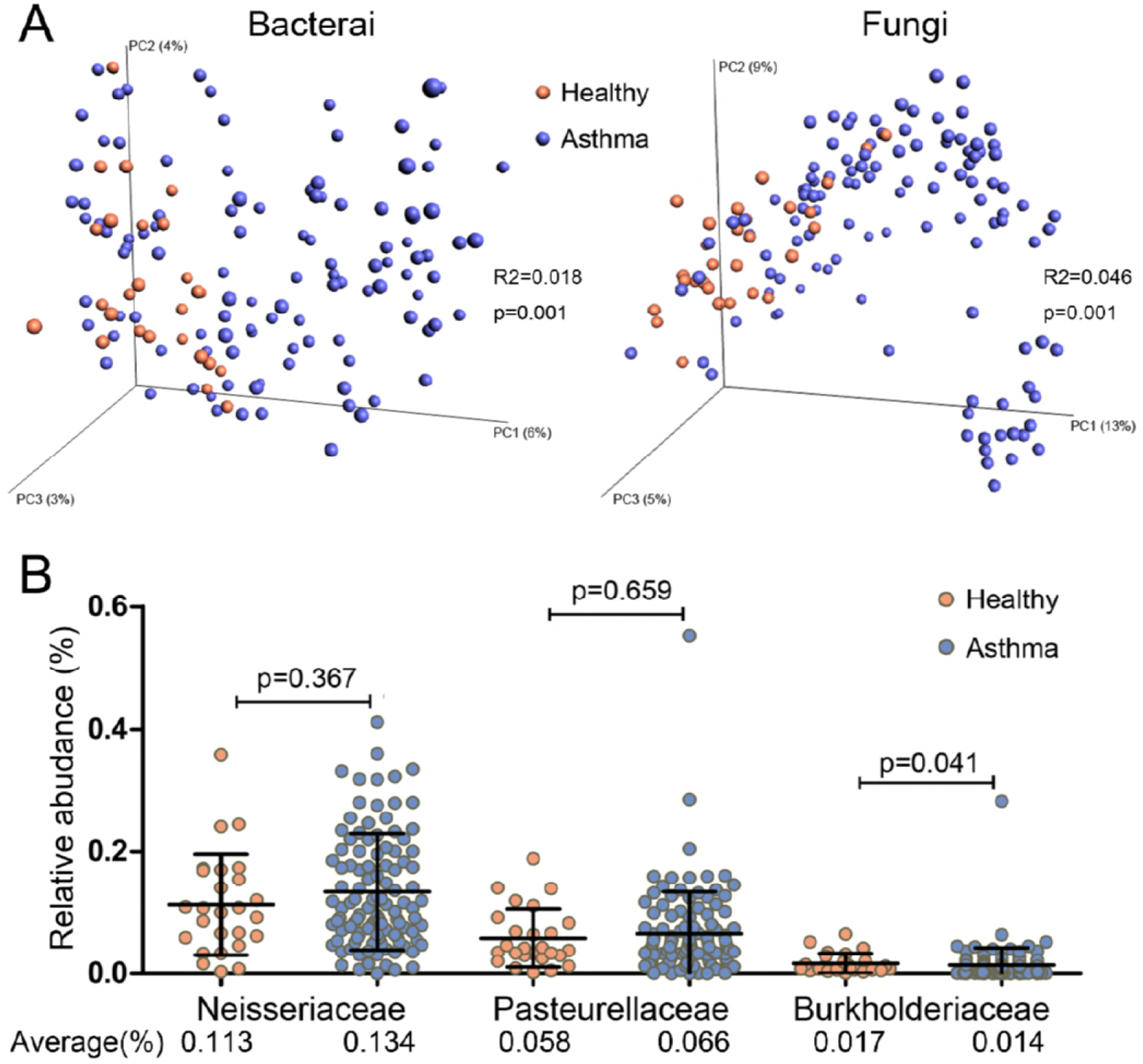
(A) Principal coordinate analysis (β-diversity) based on the unweighted_uniFrac distance of each sample of bacterial and fungal community between healthy individuals and asthma patients. (B) The relative abundances of the other families belonging to Proteobacteria compared between asthma patients and healthy individuals. P values (Wilcoxon rank-sum test) and the average relative abundance are presented.

**FIG S2.**
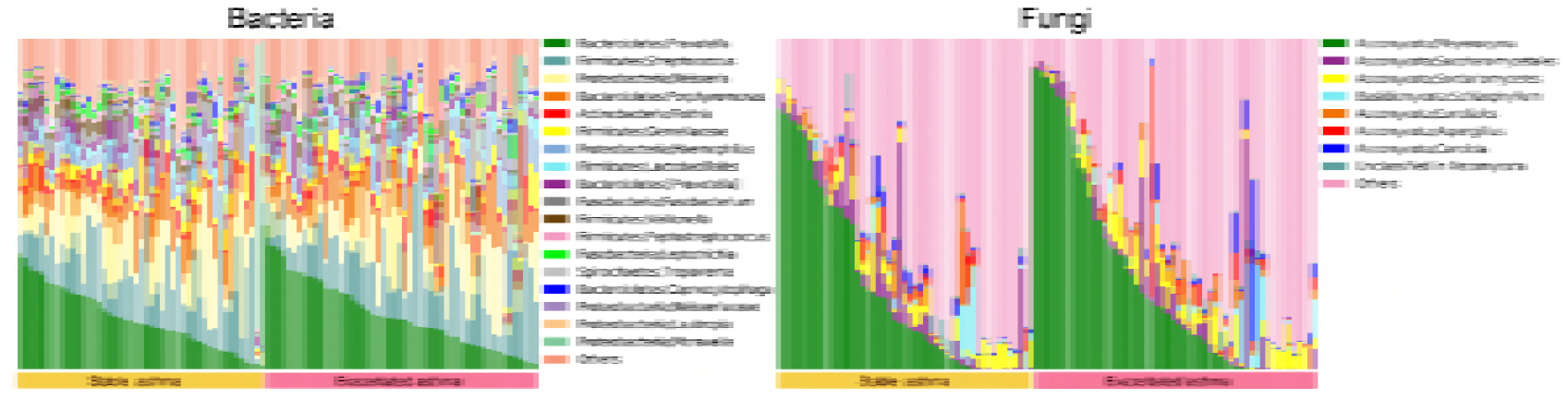
Taxonomic profiles of the most abundant airway bacterial and fungal genera in each sample from patients with stable asthma and exacerbated asthma. Genera with a relative abundance higher than 1% in any group are presented; other genera are classified as “Others”.

**FIG S3.**
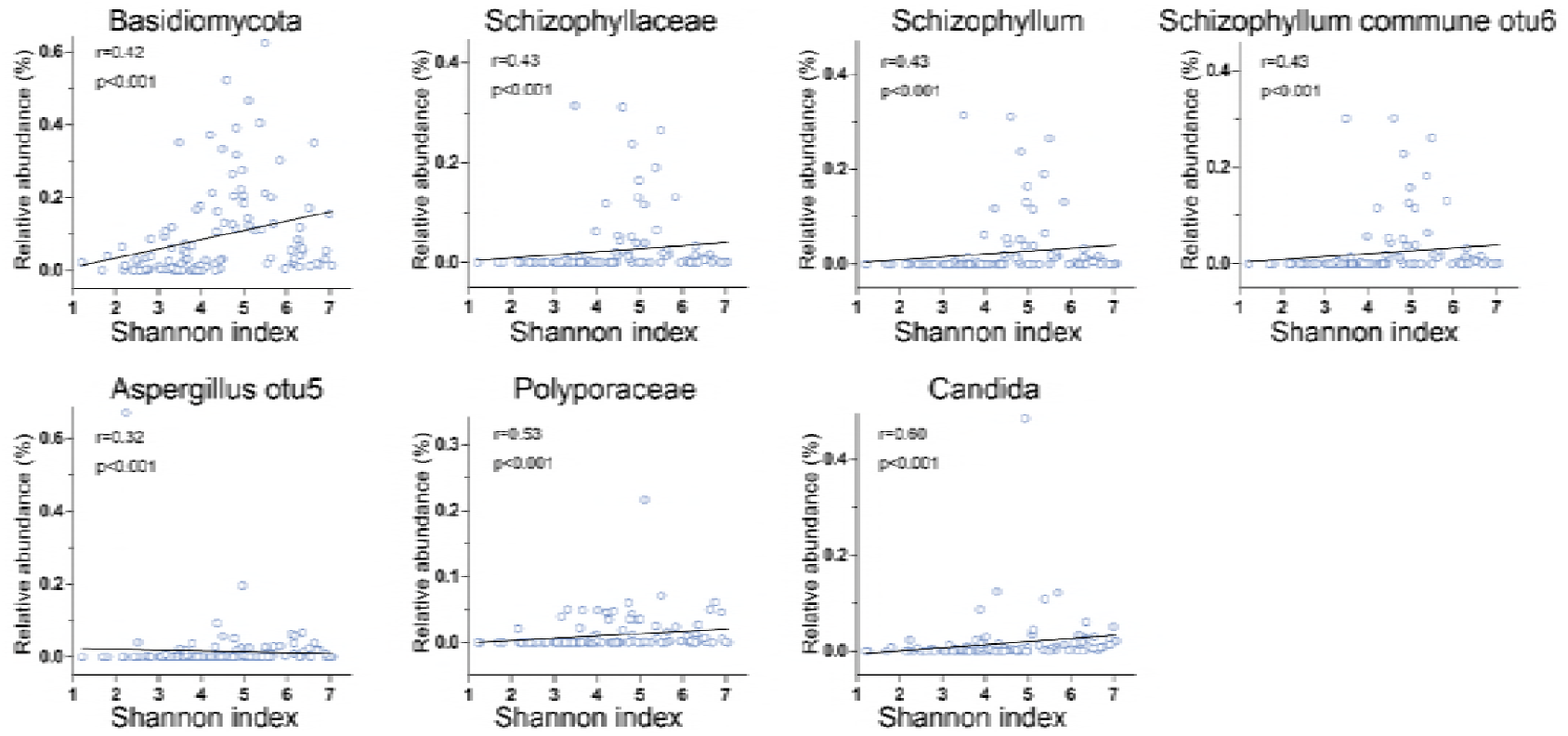
The relationship between fungal communities and fungal α-diversity. R (cut-off of over 0.3) and P values are presented using Spearman correlation analysis.

